# Does skin surface temperature variation account for Buruli ulcer lesion distribution?

**DOI:** 10.1101/760496

**Authors:** Nicola K. Sexton-Oates, Andrew J. Stewardson, Arvind Yerramilli, Paul D.R. Johnson

**Author notes:** Corresponding authors, (NKSO), (PDRJ).

## Abstract

**Background:** Buruli ulcer is a necrotising infection of skin and soft tissue caused by *Mycobacterium ulcerans* (*M. ulcerans*). Buruli ulcer most often occurs on limbs, and it is hypothesized this is explained by direct exposure to the environment. However, even on exposed areas Buruli ulcer is not randomly distributed. *M. ulcerans* prefers an in vitro temperature of 30-33°C and growth is inhibited at higher temperatures. This study investigated whether variations in skin surface temperature distribution in healthy volunteers could partly account for Buruli ulcer lesion distribution.

**Methodology/Principal Findings:** In this observational study, a thermal camera (FLIR E8) was used to measure skin surface temperature at the sternal notch and at 44 predetermined locations on the limbs of 18 human participants. Body locations of high, middle and low Buruli ulcer incidence were identified from existing density maps of lesion distribution. Skin temperature of the three incidence location groups were compared, and differences in age and sex groups were also analysed.

We found an inverse relationship between skin temperature and lesion distribution, where high incidence locations were significantly cooler and low incidence locations significantly warmer (Kruskal-Wallis test p<0.0001). Linear mixed effects regression analysis estimated that skin surface temperature accounts for 9.5% of the variance in Buruli ulcer lesion distribution (marginal R-squared = 0.095). Men had warmer upper and lower limbs than females (Mann-Whitney U test p=0.0003 and p<0.0001 respectively).

**Conclusions/Significance:** We have found an inverse relationship between skin temperature and Buruli ulcer lesion distribution, however this association is weak. Additional unknown factors are likely to be involved that explain the majority of the variation in Buruli lesion distribution.

**Author Summary:** Buruli ulcer is a destructive soft tissue infection caused by the bacterium *Mycobacterium ulcerans*. The precise mode of transmission remains unknown. One theory proposes that transmission occurs by direct contact with a contaminated environment. Lesions occur mostly on limbs, and it is hypothesized this is explained by direct exposure to the environment. However even on exposed areas, lesions are not randomly distributed. This study investigated whether skin surface temperature can partly explain Buruli ulcer lesion distribution. We measured the skin surface temperature of 18 healthy participants using a thermal camera and compared temperature distribution to the distribution of Buruli ulcer lesions investigated in a previously published study. We found that there is a negative correlation between skin temperature and Buruli ulcer lesion incidence. However, the association is weak and other factors e.g. clothing choice and insect biting patterns may explain the majority of Buruli ulcer lesion distribution.

## Introduction

Buruli ulcer is a necrotising cutaneous infection caused by the bacterium *Mycobacterium ulcerans* [1, 2]. Cases of Buruli ulcer have been reported in 33 countries, most of which are located in West and Central Africa [3], with children in these areas experiencing the majority of the disease burden [1]. Currently, there is a major outbreak occurring in south-eastern Australia on the Bellarine and Mornington peninsulas [4, 5]. Severe and/or untreated Buruli ulcer may result in contractures, deformity, permanent scarring, amputations and disabilities [6]. These can lead to social, educational and financial difficulties for those affected and their families, particularly in developing countries with limited access to modern therapy [7]. Seventy years on since the identification of *M. ulcerans* as the causative organism of Buruli ulcer, its transmission remains controversial. The disease only occurs in specific endemic locations but how exactly the infection is acquired in these regions is undetermined [5, 6].

There are several competing hypotheses concerning the transmission of *M. ulcerans*. Firstly, transmission may occur through direct contact with an environment contaminated with *M. ulcerans*, likely aided by minor cuts and abrasions sustained while working or playing outdoors [6]. Secondly, in south-eastern Australia there is increasing evidence that insects, particularly mosquitoes, may act as mechanical vectors to transmit the bacteria to humans [6]. Thirdly, *M. ulcerans* may be aerosolised from contaminated natural bodies of water, spread into the environment, then be inhaled and disseminated in the body [8]. The bacteria could then reactivate at cooler body sites [8, 9] as *M. ulcerans* prefers to grow in vitro at 30-33°C and growth is inhibited at higher temperatures [6], in a way analogous to *Mycobacterium leprae*, the causative organism of leprosy [10]. Human-to-human transmission is not thought to be of public health significance [11].

Buruli ulcer lesions are most common on limbs [1, 9, 12–16]. We postulate that skin on these areas of the body is more likely to be exposed to a contaminated environment than other areas of the body, for example the trunk. Recently, computer-generated density maps of Buruli ulcer lesion distribution have been created by analysing the locations of 649 confirmed lesions in Victoria, Australia from 1998-2015 [9]. A highly non-random distribution was found, favouring distal limbs, particularly ankles, calves and elbows. Palms of the hands and soles of the feet were rarely affected. These findings are in keeping with the mosquito vector and direct contamination hypotheses of transmission, as most lesions occurred on commonly exposed areas of the body (i.e. limbs). However, palms of the hand and soles of the feet are rarely sites of Buruli ulcer lesions. This suggests an additional factor or factors are involved in the localisation of lesions beyond just direct environmental contact. For example, trauma, insect bites, or the preference of *M. ulcerans* to grow at cooler body sites [9]. This study aimed to investigate whether skin surface temperature distribution can explain variation in Buruli ulcer incidence in different regions of the body and between different demographics (i.e. age and sex categories).

## Methods

### Study design

This was an observational study using thermal imaging to investigate skin surface temperature in volunteer participants and enable comparison to published Buruli ulcer lesion distribution data. Measurements were undertaken in a single visit per participant at the Austin Hospital between April and June 2018. Eighteen volunteer participants were included in this study, recruited in age group cohorts: ≤15 (n=2), 16-64 (n=12) and ≥65 years of age (n=4). This was to allow for comparison to published density maps of Buruli ulcer lesion distribution also categorised in these age groups. We aimed to recruit approximately 20 patients across the three age groups, based partly on time and resource availability. At the time the project was designed we were not aware of existing data on which to base a formal power calculation. We successfully recruited and studied 18 patients. We recruited a convenience sample of hospital staff, medical students, and family and friends of initial participants. Eligibility criteria included the ability to stand for 30 minutes and to be afebrile (<38°C) on the day of measurement.

### Data collection

A thermal camera (FLIR E8) was obtained to measure skin surface temperature from a distance of 30cm at the sternal notch and at 44 predetermined locations on the limbs (see Appendix 1 and 2). The thermal camera used in this study had spatial resolution identified as sufficient for the evaluation of human skin temperature [17], and the lead researcher attended a 4 hour training course provided by the manufacturer (FLIR Systems, 18/03/2018). We measured locations specifically on the limbs as these areas are commonly affected by Buruli ulcer and postulated to be commonly exposed to the environment. However within these exposed areas there is variation in lesion prevalence, and so by measuring relative temperature at different limb locations we investigated whether this variation may explain the known non-random distribution of Buruli lesions. A measurement at the sternal notch was included to enable comparison of limb measurements to the trunk and hence comparison of our findings to previous research examining limb and trunk skin temperatures. Measurements were recorded in clinic rooms at the Austin Hospital to minimise variation in room temperature and surrounding surfaces, as these can affect skin surface temperature and thermal camera measurements. Two temperatures were recorded for each location, measured approximately 15 minutes apart. Thermal images of upper and lower limbs were also recorded from 1.5 and 3 metres.

Participants rested in the clinic room for approximately 10 minutes prior to measurement to minimise the effect of prior physical activity on skin surface temperature distribution. Participants also completed a questionnaire regarding age, sex and a number of medical conditions/medications known to affect skin surface temperature (see Appendix 3). Room temperature was recorded using Aqua Systems ‘Wooden Wall Thermometer’. Core body temperature was measured using a temporal artery thermometer (Exergen TAT-5000) to ensure participants were afebrile. Core body temperature, sternal notch and left cubital fossa temperature measurements of a control, the investigator, were recorded at each session to examine the consistency of skin surface temperature measurements over time and with varying room and core body temperatures.

### Categorisation of body locations of high, medium and low Buruli ulcer incidence

Previously published density maps of Buruli ulcer lesion distribution (Fig 1) were created using a 15-layer colour ramp from green (lowest density, 1/15) to red (highest density, 15/15) as per the first author of the publication. Using these published maps, density gradations were assigned to the body locations investigated in this study (Table 1).

**Fig 1.**
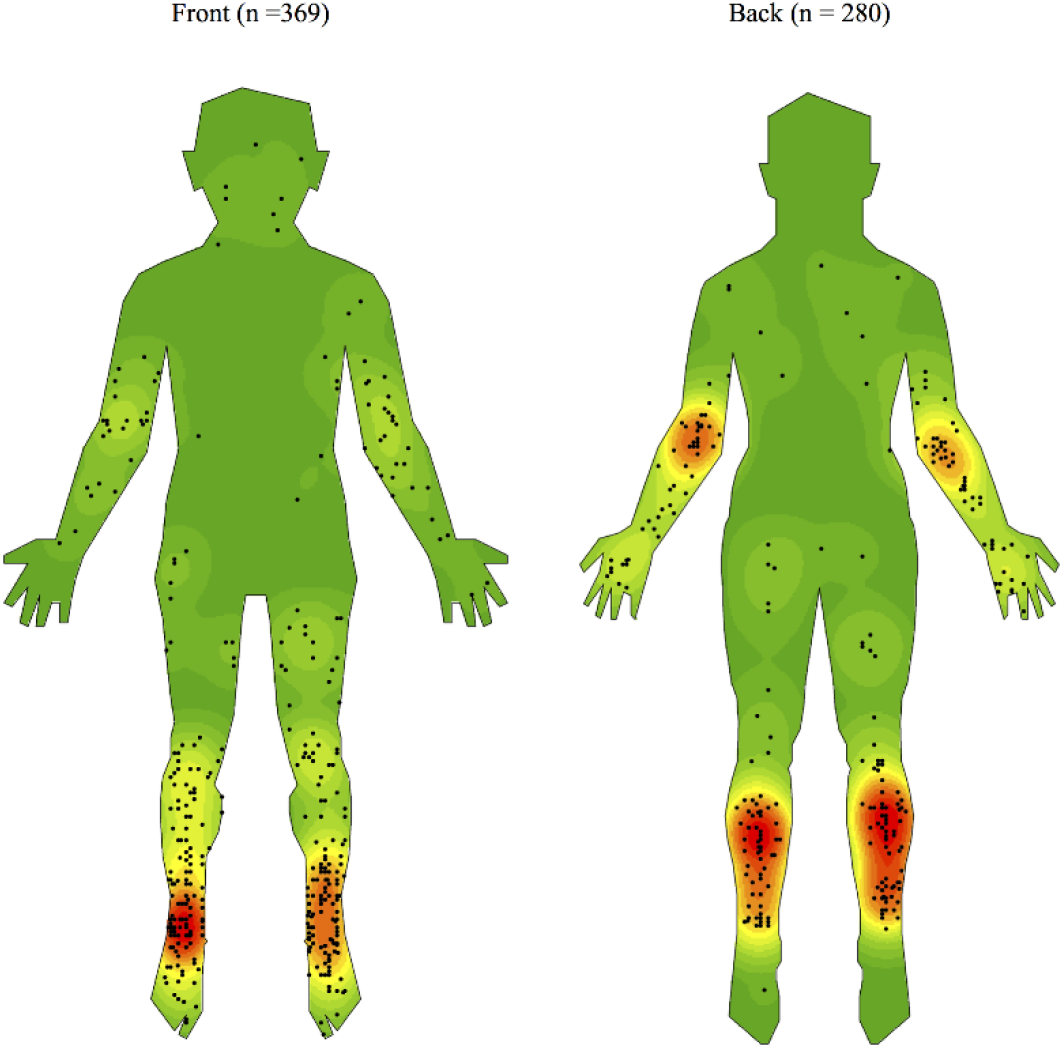
Density map of the distribution of Buruli ulcer lesions on front and back of human body templates generated using ArcGIS software version 10.3.1. [9]

**Table 1.** Body locations with corresponding Buruli ulcer lesion density gradations, categorised into high, medium and low Buruli ulcer incidence groups.

| Location | Density gradation |
| --- | --- |
| **High Buruli ulcer incidence** |  |
| Posterior upper calf | 8-14 |
| Posterior mid calf | 14-15 |
| Posterior lower calf | 7-12 |
| Anterior lower shin | 8-15 |
| Medial malleolus | 10-13 |
| Lateral malleolus | 10-15 |
| Elbow | 10-12 |
| **Medium Buruli ulcer incidence** |  |
| Dorsum of hand | 5-7 |
| Posterior mid forearm | 6-7 |
| Dorsum of foot | 7-9 |
| Anterior upper shin | 5-7 |
| Anterior mid shin | 7-9 |
| Anterior knee | 6-7 |
| **Low Buruli ulcer incidence** |  |
| Posterior mid arm | 3-4 |
| Posterior mid thigh | 2-3 |
| Popliteal fossa | 3-5 |
| Sole of foot | 1 |
| Sternal notch | 1 |
| Anterior mid thigh | 1-3 |
| Central palm | 1-2 |
| Anterior mid forearm | 2-3 |
| Cubital fossa | 5 |
| Anterior mid arm | 3-4 |

Body locations of high Buruli ulcer incidence for this study have been defined as areas of lesion density in the highest third of density gradations, corresponding to gradations ≥ 11/15. Body locations of medium Buruli ulcer incidence have been defined as areas of lesion density in the middle third of density gradations, corresponding to gradations 6-10/15. Body locations of low Buruli ulcer incidence have been defined as areas with density gradation 1-5/15. If a location had a range of density gradations, the highest gradation was used to determine Buruli ulcer incidence category.

### Statistical analysis

The mean skin surface temperature of each body location was compared to the mean of all other locations combined using Mann-Whitney *U* tests. The median temperatures of the three Buruli ulcer incidence groups were compared using a Kruskal-Wallis test. Differences in age and sex groups were also analysed using Kruskal-Wallis and Mann-Whitney *U* tests.

Additionally, we built a mixed-effects linear regression model using to quantify the association between temperature and incidence of Buruli ulcer in each body location. We accounted for the repeated temperature measurements by including random effects for participant (random intercept and slope) and for the potential impact of age by including a random slope for age.

The regression model was built using R, version 3.4.4 (R Foundation for Statistical Computing, Vienna, Austria). All other analyses were performed using GraphPad Prism^®^ version 7.04.

### Ethical statement

This study was approved by the Austin Health Human Research Ethics Committee. Reference number: HREC/17/Austin/578. Written consent was obtained from each participant.

## Results

### Participant cohort analysis

Eighteen participants were included. The mean age was 38.7 years (range = 11.8-77.3, IQR = 24.7-64.4). Nine participants were male (50%) and 9 female (50%). In the ≤ 15 years age group (n=2), there were 2 (100%) female participants. In the 16-64 years age group (n=12), there were 6 (50%) females and 6 (50%) males. In the ≥ 65 years age group (n=4), there was 1 (25%) female and 3 (75%) males.

Of the 18 participants, 4 (22%) reported having experienced Chilblains, 1 (6%) peripheral vascular disease, 1 (6%) suspected Raynaud’s phenomenon, 1 (6%) low-functioning thyroid on thyroxine with normal TSH (thyroid-stimulating hormone) levels, and 1 (6%) taking a blood pressure medication (Irbesartan). No participants reported having diabetes for > 5 years, a high-functioning thyroid, neuropathy, sunburn or taking migraine medication.

### Grouping of left and right measurements

With the exception of the sternal notch, measurements were recorded on both sides of the body for each location, e.g. left anterior knee and right anterior knee. As there was no significant difference between left and right measurements by Mann-Whitney *U* test (p=0.4844), these groups were combined to give 23 body locations for reporting of mean skin surface temperature and further analysis.

### Mean skin surface temperature in high, medium and low Buruli ulcer incidence locations

Participant skin surface temperature measurements ranged from 22.6 to 35.3°C, with a mean of 30.1°C (Table 2). Skin surface temperature data was not normally distributed (D’Agostino-Pearson normality test p<0.0001). Cubital fossa was the location of highest mean skin surface temperature (33.2°C) and sole of foot the location of lowest mean skin surface temperature (27.7°C) (Table 2 and Fig 2). Overall, the three incidence groups were found to have significantly different median temperatures (Kruskal-Wallis test p<0.0001). The high incidence group was the coolest, the low incidence group the warmest, and the medium incidence group fell in between (median 29.3, 31.1 and 30.6°C respectively). This is reflected in figures 2 and 3, which show a visually-apparent negative correlation between Buruli ulcer incidence and mean skin surface temperature. This is supported by linear mixed effects regression analysis, which estimates that skin surface temperature accounts for 9.5% of the variance in Buruli ulcer lesion distribution (marginal R-squared = 0.095). Additionally, for each one degree (Celsius) increase in the temperature of a body location there is a 0.59 (95% CI, 0.78–0.41) reduction in incidence category of that part of the body.

**Fig 2.**
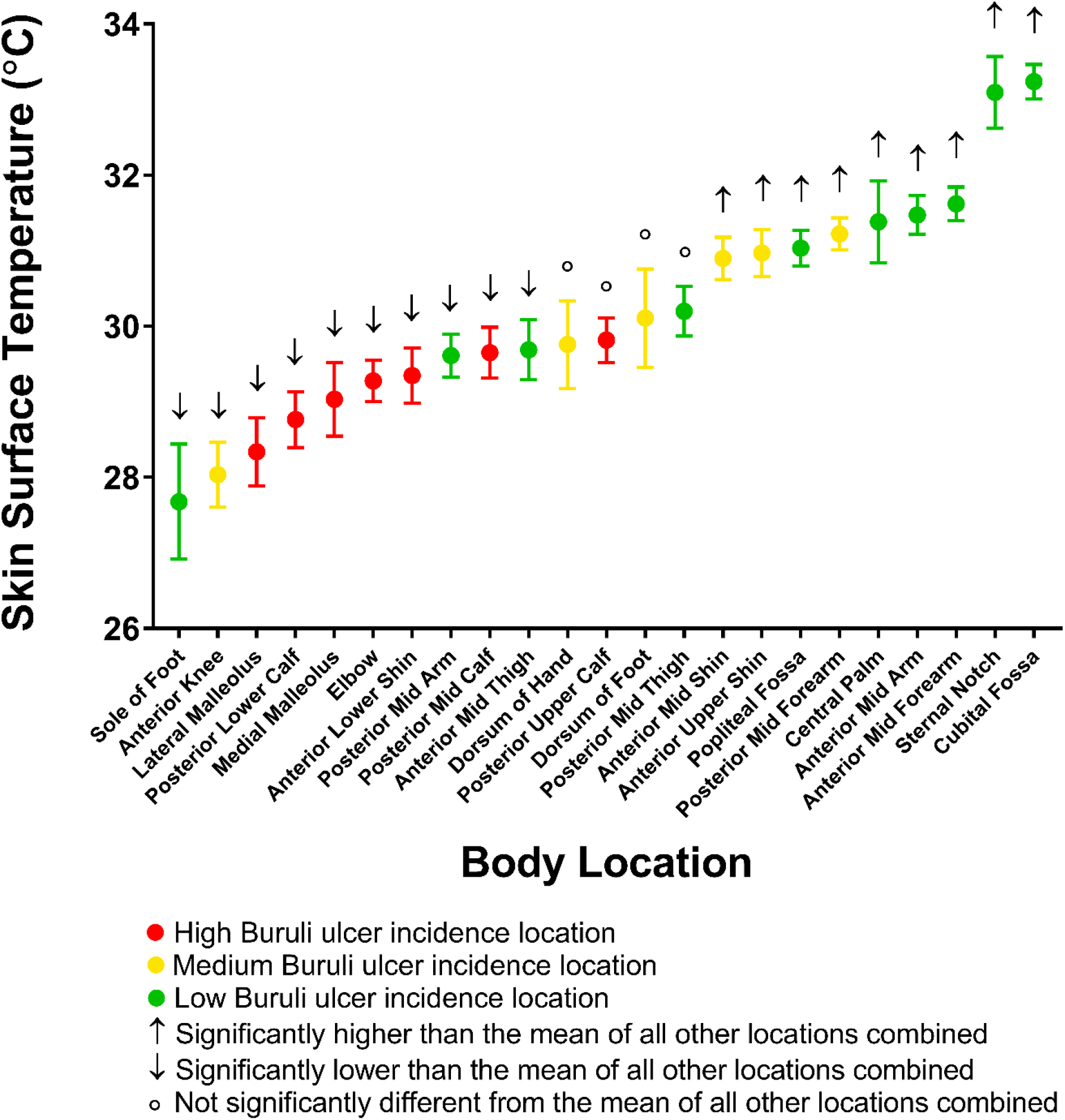
Mean skin surface temperature with 95% CI for each body location. Results of comparison to mean skin surface temperature of all other locations combined by Mann-Whitney *U* test are also shown (p values from left to right: p<0.0001, p<0.0001, p<0.0001, p<0.0001, p<0.0001, p<0.0001, p=0.0003, p=0.0033, p=0.0119, p=0.0293, p=0.125, p=0.0601, p=0.8182, p=0.9846, p=0.0008, p=0.0005, p<0.0001, p<0.0001, p<0.0001, p<0.0001, p<0.0001, p<0.0001, p<0.0001)

**Fig 3.**
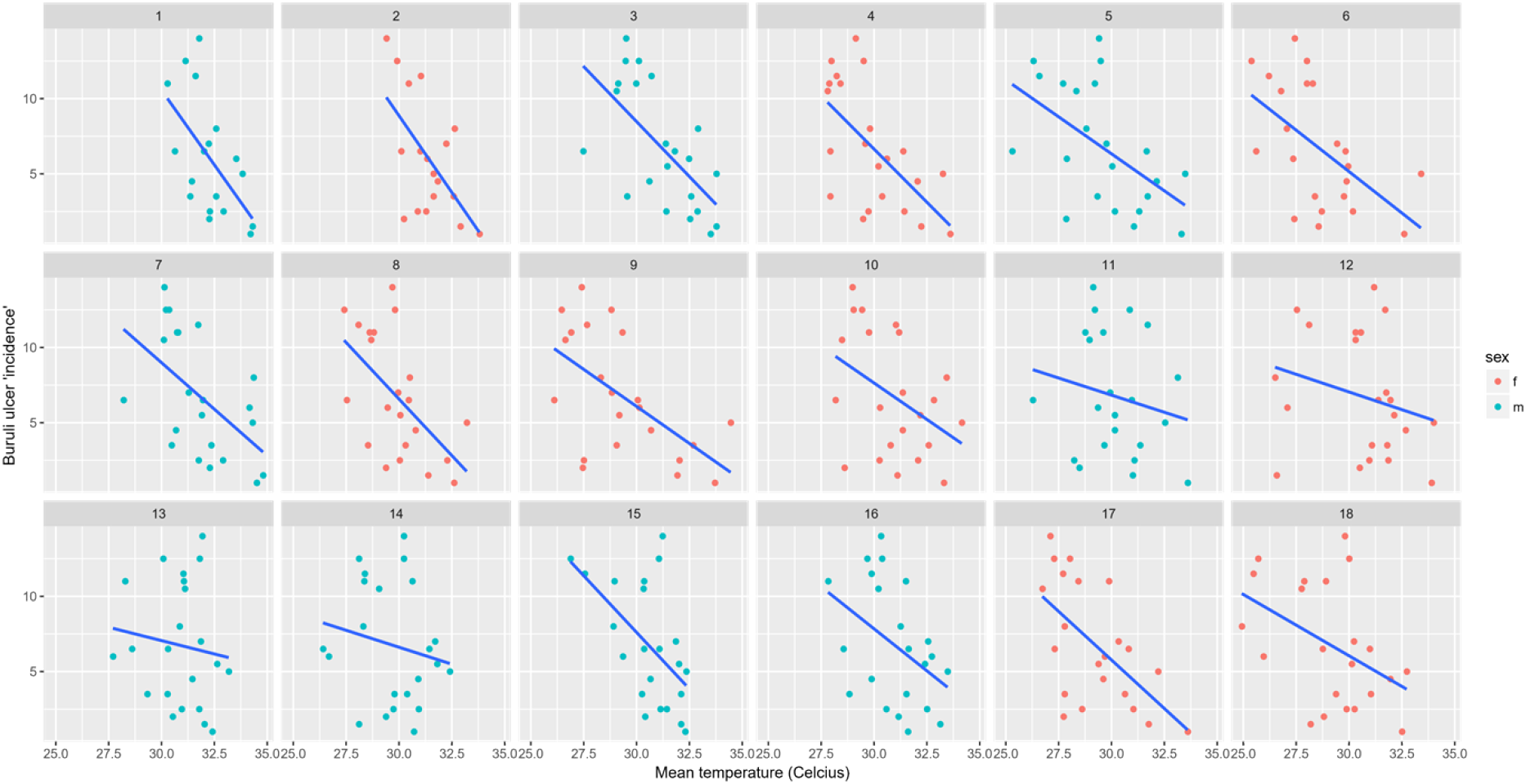
Scatterplot of mean temperature and Buruli ulcer incidence gradation for each body location, overlayed by a line of best fit. Each participant’s is data shown separately. The data points for the location ‘sole of foot’ have been excluded.

**Table 2.** Mean and range of temperature recordings.

| **Measurement** | **Mean (°C)** | **Range (°C)** |
| --- | --- | --- |
| **Anterior and Posterior Locations** | 30.1 | 22.6-35.3 |
| **Anterior Locations** |  |  |
| Sternal notch | 33.1 | 30.7-34.5 |
| Central palm | 31.4 | 25.4-35.2 |
| Mid forearm | 31.6 | 30.0-33.9 |
| Cubital fossa | 33.2 | 31.1-35.3 |
| Mid arm | 31.5 | 29.2-33.4 |
| Dorsum of foot | 30.1 | 23.8-35.0 |
| Medial malleolus | 29.0 | 24.7-32.2 |
| Lower shin | 29.4 | 26.6-32.0 |
| Mid shin | 30.9 | 28.5-32.9 |
| Upper shin | 31.0 | 27.5-32.9 |
| Knee | 28.0 | 25.0-32.1 |
| Mid thigh | 29.7 | 26.7-32.8 |
| **Posterior Locations** |  |  |
| Dorsum of hand | 29.8 | 25.7-34.7 |
| Mid forearm | 31.2 | 29.0-33.6 |
| Elbow | 29.3 | 27.0-32.0 |
| Mid arm | 29.6 | 27.4-32.2 |
| Sole of foot | 27.7 | 22.6-32.7 |
| Lateral malleolus | 28.3 | 24.2-32.0 |
| Lower calf | 28.8 | 25.8-31.6 |
| Mid calf | 29.7 | 26.2-32.4 |
| Upper calf | 29.8 | 27.1-32.5 |
| Popliteal fossa | 31.0 | 28.1-32.9 |
| Mid thigh | 30.2 | 26.8-32.8 |
| **Additional Measurements** |  |  |
| Control sternal notch | 33.9 | 32.5-35.3 |
| Control left cubital fossa | 33.7 | 32.5-34.7 |
| Control core body temperature | 36.6 | 36.3-36.9 |
| Room temperature | 21.1 | 20.0-25.0 |
| Participant core body temperature | 36.5 | 36.0-37.0 |

With respect to the 7 high Buruli ulcer incidence locations, 6 (86%) had mean skin surface temperature significantly lower than the mean of all other locations combined when testing with Mann-Whitney U tests (Fig 2). These locations were elbow, anterior lower shin, lateral malleolus, medial malleolus, posterior lower calf and posterior mid calf. One (14%) high Buruli ulcer incidence location, posterior upper calf, did not have mean temperature significantly different from the mean of all other locations combined. No high Buruli ulcer incidence locations had mean temperatures significantly higher than the mean of all other locations combined. Notably, two low incidence locations were also within the same temperature range as high lesion density locations; anterior mid thigh and posterior mid arm.

### Comparison by age group and sex

Analysis of skin surface temperatures by sex and age group are shown in Table 3. All male and all female measurements were compared using a Mann-Whitney *U* test.

**Table 3.** Comparison of mean skin surface temperature by age group and sex.

| Location | Mean skin temperature ($^{\circ}\text{C}$) | | | p value |
| --- | --- | --- | --- | --- |
|  | Male | Female |  |  |
| All locations | 30.6 | 29.6 | | $<0.0001$ (Mann-Whitney U) |
| Upper limb | 31.5 | 30.8 |  | 0.0003 (Mann-Whitney U) |
| Lower limb | 30.2 | 28.9 | | $<0.0001$ (Mann-Whitney U) |
| | $\leq 15$ years | 16-64 years | $\geq 65$ years | |
| All locations | 28.1 | 30.3 | 30.3 | $<0.0001$ (Kruskal-Wallis) |
| | $<65$ years | | $\geq 65$ years | |
| Mid arm | 30.6 |  | 30.4 | 0.433 (Mann-Whitney U) |

The male group had a significantly higher skin surface temperature overall than the female group (p<0.0001), with a mean of 30.6°C compared to 29.6°C respectively. When comparing all upper limb measurements, males had a significantly higher mean skin surface temperature than females (31.5 and 30.8°C respectively, p=0.0003). When comparing all lower limb measurements, males also had a significantly higher mean skin surface temperature than females (30.2 and 28.9°C respectively, p<0.0001).

The skin surface temperature measurements of the three age groups were compared using a Kruskal-Wallis test and found to be significantly different (p<0.0001). The ≤15 age group had an overall mean skin surface temperature of 28.1°C (range 23-33.6), the 16-64 age group 30.3°C (range 24-35.3) and the ≥ 65 age group 30.3°C (range 22.6-34.7). However, these data should be considered with caution as we were only able to recruit 2 participants for the ≤15 age group.

To enable comparison to previously published work, anterior and posterior mid arm temperatures combined were compared between those <65 years and those ≥65 years. No significant difference was found by Mann-Whitney *U* test (p=0.433, means 30.59 and 30.37°C respectively).

### Control measurements

Measurements of the constant control’s core body temperature, sternal notch and left cubital fossa skin temperature varied throughout the data collection period (see Table 1). Sternal notch ranged from 32.5-35.3°C, left cubital fossa 32.5-34.7°C, and core body temperature 36.3-36.9°C. The constant control was taking the oral contraceptive pill during the measurement period and so menstrual cycle hormonal changes were not expected to affect core body temperature.

## Discussion

We have found that overall, the three Buruli ulcer incidence location groups derived from previously published work had significantly different median skin surface temperatures. The highest incidence group had the coolest median temperature, the lowest incidence group had the highest median temperature, and the middle incidence group fell in between. In addition, the linear mixed effects regression analysis estimated that skin surface temperature accounts for 9.5% of the variance in Buruli ulcer lesion distribution (marginal R-squared = 0.095). This generally supports a previously stated hypothesis that Buruli ulcer lesions occur preferentially on areas of relatively lower skin temperature. However, two low incidence areas (anterior mid thigh and posterior mid arm) were within the same skin surface temperature range as the high incidence group. Additionally the coolest region of all, sole of foot, is rarely affected by Buruli ulcer.

Buruli ulcer lesion distribution has been found to differ between men and women, and between age groups. For example, men have been found to be more likely than females to have lesions on upper limbs, and less likely to have lesions on lower limbs [9, 15]. Those ≥65 years of age have been found to be less likely to have a lesion on the arm and shoulder than those <65 [9]. We investigated whether these differences between age and sex correlated with differences in skin surface temperature distribution. We found that men had significantly higher skin surface temperatures than females for both upper and lower limbs. There was no significant difference in mid arm measurements found between those ≥65 and those >65. As such, we conclude that differences in skin surface temperature distribution found in this study do not account for differences in Buruli ulcer lesion distribution between age and sex groups.

Our findings add some support to the aerosol-dissemination and reactivation hypothesis of transmission as we found a negative correlation between skin surface temperature and Buruli ulcer lesion distribution. However, the association is weak and there are important exceptions (e.g. sole of foot) where the relationship breaks down. With respect to the mosquito vector hypothesis, skin temperature may influence mosquito biting patterns and hence Buruli ulcer lesion distribution. A future direction of research may be the direct investigation of the biting patterns of mosquitoes, particularly of species hypothesised to mechanically transmit *M. ulcerans* in south-eastern Australia.

To our knowledge, the skin surface temperatures of high, medium and low Buruli ulcer incidence lesion locations have not been previously studied. Several studies have examined skin temperature distribution more generally and found that the trunk is warmer than limbs, and proximal areas of limbs warmer than distal areas [18, 19]. The results from this study are consistent with these observations, finding that the sternal notch on the trunk had a mean skin surface temperature higher than 21 of 22 measured limb locations. In addition, the mean foot measurements were colder than mean mid thigh measurements, and the mean dorsal hand temperature was colder than mean posterior mid arm. In contrast, the mean temperature for palm of hand was warmer than that of the anterior mid arm.

The validity of thermal camera measurements and control of the factors which may influence them are important to consider. Thermal cameras have been used to record skin surface temperature in both disease and non-disease states [20]. We have used a thermal camera identified as appropriate for clinical research in humans [17], and believe our method of measurement (on-the-spot readings from a distance of 30cm, as opposed to taking measurements from a thermal image at a greater distance using a software program) optimises the validity and consistency of temperature readings. This conclusion draws from the training course run by FLIR Systems. Surfaces within the room and properties of the measured surface (in this case, skin) affect thermal camera measurements [21]. We controlled these variables by adjusting the thermal camera emissivity setting to 0.98, appropriate for human skin [21], and conducting the measurements in clinic rooms at the Austin hospital containing similar surfaces.

Limitations of this study include the fact that specific points were used to represent temperature for a larger area, e.g. olecranon fossa for elbow. Within a defined area there are often multiple smaller areas of differing temperature (see Appendix 4), and as such the selected measurement points may not accurately represent the temperature of the larger area. In addition, the number of measurement locations may limit the generalisability of our findings to the whole body. A further limitation is the variance in room temperature (20-25°C) as this may have affected skin surface temperature and has not been taken into account in our analyses. The variation of skin surface temperature of the control was 2.8°C for the sternal notch and 2.2°C for the cubital fossa, and may be due to genuine fluctuation in skin surface temperature or inconsistent measurement. A further limitation is that this study included a small number of participants with conditions likely to affect skin surface temperature e.g. peripheral vascular disease [17]. Participants rested for at least 10 minutes prior to measurements to minimise the effect of prior physical activity, however other factors that may influence skin surface temperature and its distribution (e.g. emotional state) are difficult to ascertain and were not controlled for. Lastly, the non-Gaussian distribution of the data and correlation between adjacent regions of the body may limit the appropriateness of the linear regression model.

In conclusion, we have found that there is an inverse relationship between skin surface temperature in healthy volunteers and previously published Buruli ulcer lesion distribution. However relative skin temperature appears to be only weakly associated with Buruli lesion distribution, meaning that more than 90% of the clinically observed non-random distribution is likely to be explained by other factors such as clothing choice, skin trauma and targeting behaviour by insects.

## Acknowledgements

We would like to acknowledge the participants of this study and thank them for volunteering their time.

**Supporting Information Captions**

**S1 Appendix.** Skin temperature measurement procedure. This image displays the thermal camera screen during measurement. The crosshairs (labelled Sp1) indicate the point of measurement, and the temperature reading in the top left hand corner of the image shows the measured temperature of that area.

**S2 Appendix.** Skin temperature measurement locations.

**S3 Appendix.** Study questionnaire.

**S4 Appendix.** Selection of participant thermographs. a) Anterior lower limbs, b) Anterior upper limbs, c) Posterior lower limbs, d) Posterior upper limbs

**S5 Appendix. STROBE Checklist.**

